# Midgut damage triggers thromboxane A_2_–dependent hemocyte recruitment in *Culex pipiens molestus*

**DOI:** 10.64898/2026.07.02.735721

**Authors:** Dawon Lee, Du-Yeol Choi, Dongjun Kang, Singeun Oh, Arwa Shatta, Myung-hee Yi, Jun Ho Choi, Yun Soo Jang, Chaerin Park, In-Yong Lee, Ju Yeong Kim

## Abstract

Mosquitoes transmit diverse pathogens through repeated blood feeding, a process that subjects the midgut to significant mechanical stress and cellular damage. While hemocyte association with the midgut is known to occur following injury, the mechanisms promoting their recruitment remain poorly defined. Given the conserved roles of eicosanoid signaling in injury responses, we hypothesized that thromboxane A_2_ (TXA_2_) mediates hemocyte recruitment to the damaged midgut. Here, we demonstrate that chemically induced midgut damage significantly increases both the number of hemocytes attached to the midgut and those in circulation. Pharmacological inhibition of cyclooxygenase suppresses hemocyte recruitment whereas supplementation with a stable TXA_2_ analog restores the response, indicating that TXA_2_ signaling is required for this process. To identify candidate enzymes involved in TXA_2_ biosynthesis, we performed *in silico* docking analyses and identified two cytochrome P450 (CYP) candidates. Among these, *CYP6D3* was shown to be strongly upregulated in hemocyte populations following midgut damage. RNA interference-mediated knockdown of *CYP6D3* significantly reduced both hemocyte recruitment to the midgut and systemic TXB_2_ levels, supporting its role in the eicosanoid-mediated immune response. Together, our findings demonstrate that TXA_2_ signaling drives hemocyte recruitment to the damaged mosquito midgut and suggest a conserved lipid-mediated mechanism underlying insect tissue-associated immune responses that may influence vector competence.

## 1. Introduction

Vector-borne diseases, especially mosquito-borne diseases (MBDs), account for more than 700 million incidences of infection and an estimated one million deaths annually worldwide (Chandra and Burman, 2024). *Culex spp*. transmit pathogens of clinical importance such as West Nile virus and Japanese encephalitis virus and are also involved in transmission of veterinary diseases such as avian malaria and filariasis. Recent studies have also reported varying degrees of vector competence for emerging viral pathogens such as Rift Valley Fever virus, Usutu virus, and St. Louis encephalitis virus (Schulz and Becker, 2018).

For transmission of MBDs, the mosquito must first acquire pathogens from an infected host through a blood meal. After ingestion, the pathogen crosses the midgut barrier and infects tissues that support its replication. Following egress from infected tissues, the pathogen enters the hemolymph in a process known as dissemination. These events occur during the extrinsic incubation period (EIP), the time required for a pathogen to develop within the mosquito before transmission becomes possible (Franz et al., 2015). The EIP is a crucial stage in MBD transmission and a stage unique from others in that it requires pathogen survival against the mosquito immune system (Prince et al., 2023). Because pathogen dissemination begins early in the EIP, immune responses triggered immediately after midgut damage must be precisely timed and regulated to influence transmission success. The final stage of transmission is the migration of the pathogen to the salivary glands where it can be transmitted again to a new host during a subsequent blood meal.

The mosquito midgut serves as the primary physical and physiological barrier against dissemination and propagation of pathogens within the mosquito (Lewis et al., 2023). During blood feeding, midgut barrier becomes distended and damaged. In Anopheles spp., the damaged midgut is known to undergo regeneration by the recruitment of hemocytes (Cardoso-Jaime and Dimopoulos, 2025). Hemocytes prevent further infection and promote wound healing, first by aggregating to sites of damage and subsequently releasing downstream signals that mediate coordination of immune responses such as melanization and phagocytosis (Hillyer and Strand, 2014). Although hemocytes contribute to midgut repair and help limit pathogen dissemination, the molecular regulators that govern hemocyte recruitment in *Culex* mosquitoes remain poorly understood. Most mechanistic insights have been derived from *Anopheles* or *Aedes* species, leaving key immune signaling pathways in *Culex* largely unexplored.

Eicosanoids are well-characterized signaling lipids in mammals, where they play key roles in vasoconstriction and platelet aggregation (Sheppe and Edelmann, 2021). They are also present in insects, where they have been implicated in development, immunity, and reproduction (Stanley and Kim, 2019). yet their specific types and immunological functions remain poorly understood (Kim and Stanley, 2021). One of the few well-characterized examples is Prostaglandin E_2_ (PGE_2_), which facilitates hemocyte recruitment and immune activation in *Anopheles* spp. by modulating hemocyte composition, increasing the proportion of granulocytes during bacterial infection (Barletta et al., 2019). Despite this, the functions of many eicosanoids in insect immunity remain largely uncharacterized.

Among these, TXA_2_ has been implicated in early immune responses. TXA_2_ is highly unstable and rapidly hydrolyzes to TXB_2_ within seconds to minutes, making TXB_2_ a common indicator used in place to measure TXA_2_ production. In insects, TXB_2_ was identified in *Spodoptera exigua* using liquid chromatography, and the biosynthetic potential for TXA_2_ production was suggested by RNA interference (RNAi) experiments (Roy et al., 2021). TXA_2_ is associated with rapid cell aggregation and barrier regulation, indicating a role in coordinating early hemocyte responses following tissue damage. It signals through a G protein-coupled receptor, triggering phospholipase C–dependent calcium mobilization and downstream cellular responses, supporting its potential role in hemocyte activation during early gut damage.

In this study, we investigated the role of TXA_2_ in hemocyte mobilization during the early stages of midgut damage in mosquitoes. Our findings provide insight into early hemocyte responses associated with midgut tissue damage.

## 2. Materials and methods

### 2.1. Mosquito rearing

A laboratory strain of female mosquitoes, *Culex pipiens molestus*, were used for this study. Mosquitoes were reared in laboratory conditions at 28^°^C and 70% relative humidity with a 16/8-hour day/night cycle. Larvae were fed with a sterilized mixture of fish food and heat-killed yeast. Adults were supplied with a 7% sterilized sucrose solution via a cotton pad and were allowed to feed *ad libitum*. Mosquitoes used for experiments were collected as pupae and allowed to eclose as adults in paper cups covered with gauze cloth under the same conditions. Mosquitoes used for assays were all allowed to mature between 4-6 days prior to experimentation.

### 2.2. Chemical treatment of the mosquito midgut

Mosquitoes were starved for a minimum of 1h before SDS treatment. A sterilized cotton pad soaked in 10 ml of 7% sucrose for controls, or a 10 ml mixture of 2% Sodium Dodecyl Sulfate (SDS) in 7% sucrose for treatment was placed on the paper cup containing 20-40 female mosquitoes. The mosquitoes were allowed to feed *ad libitum* for 16-24 h at which point the mosquitoes were cold-anesthetized and moved to a new cup to prevent further exposure to treatment solutions. To confirm sucrose feeding, adult female mosquitoes were deprived of sucrose for 4 h and then allowed to feed *ad libitum* for 16 h on 7% (w/v) sucrose containing 2.5% (w/v) Erioglaucine disodium (FD&C Blue No. 1; Sigma-Aldrich, Burlington MA, USA). Following the feeding period, mosquitoes with visibly blue abdomens were considered to have successfully ingested the sucrose solution. For blood-feeding, mosquitoes were also starved and then provided defibrinated sheep blood containing anticoagulant mixed with 7% sucrose at a 2:3 ratio. Preliminary feeding assays showed poor voluntary feeding on higher blood concentrations or undiluted blood under these experimental conditions; therefore, the blood-sucrose mixture was used to achieve consistent feeding. Mosquitoes were allowed to feed in the dark *ad libitum* for 16 h.

### 2.3. Midgut treatment with inhibitors and recovery agents

Inhibition assays were conducted by injecting cold-anesthetized mosquitoes with 100 nL of 1% Dimethyl sulfoxide (DMSO) for controls and 100 ng/µl acetylsalicylic acid (ASA, aspirin) dissolved in 1% DMSO for treatment. *In vivo* recovery experiments were conducted by 100 nL injection of 1% DMSO for controls and the stabilized TXA_2_ analog, Carbocyclic TXA_2_ (CTA2; Cayman Chemicals, Ann Arbor, MI, USA) or prostaglandin E_2_ (PGE_2_; Cayman Chemicals, Ann Arbor, MI, USA) for treatments at concentrations ranging from 100 ng/µl to 1000 ng/µl 16 h following inhibitor treatment. Mosquitoes were given sucrose on a cotton pad and kept in paper cups at RT between treatments.

### 2.4. Isolation of mosquito midguts

A microinjector (UltraMicroPump3 and SMARTouch Controller, Sarasota, FL, USA) was used to inject mosquitoes with 20% formaldehyde, or mosquitoes were cold anesthetized by placing the cups on ice and subsequently transferred to a chilled plate. A stereomicroscope (Zeiss Stemi DV4 Stereo Microscope; Zeiss, Jena, Germany) was used to dissect and isolate the midgut. The head and wings were removed by forceps, and the posterior abdominal segments were pulled to expose the midgut. Other appendages such as the crop and foregut were removed for precise imaging and consistency.

### 2.5. Mosquito hemocyte counting and staining

Hemolymph collection was performed using a modified protocol (Qayum and Telang, 2011). Mosquitoes were cold-anesthetized and injected with 4,500 nL of anticoagulant RPMI (20 mM sodium citrate in RPMI 1640). After a 20-min incubation at room temperature, five female mosquitoes were reinjected with RPMI, and the last two abdominal segments were excised. Hemolymph (4.5 µL) was collected using a glass capillary and transferred to a low protein-binding microcentrifuge tube (Thermo Fisher Scientific, Waltham, MA, USA) on ice. Diff-Quik staining was performed according to the manufacturer’s instructions (Sysmex Corporation, Kobe, Japan) to distinguish different hemocyte types. Stained hemocytes were observed under a stereomicroscope (Zeiss Stemi DV4; Zeiss, Jena, Germany) and classified based on representative morphological characteristics.

### 2.6. Fixation and fluorescent staining

Staining with fluorescent dyes was performed to visualize the structure and hemocytes attached to the mosquito midgut. For visualization of hemocytes, cold-anesthetized mosquitoes were injected with 200 nL of Vibrant CM-Dil Cell-Labeling Solution (Thermo Fisher Scientific, Waltham, MA, USA) at 1:10 dilution and incubated at RT for 30 mins. Following fixation by 200 nL of 20% formaldehyde, dissection was performed in 1× PBS and midguts were transferred to 4% formaldehyde for additional fixation. Midguts were washed 3 times by 1× PBS and blocking and permeabilization were performed by 30 mins incubation in PBSBT (1× PBS, 1% BSA, 0.1% Triton X-100) for blocking. To visualize midgut structures, additional staining was performed by incubating the midgut in a mixture of 1:500 Hoechst 33342 (Invitrogen, Carlsbad, CA, USA) solution and 1:400 of the Alexa Fluor™ 488-conjugated phalloidin (Invitrogen, Carlsbad, CA, USA) solution for 15 mins. The samples were then washed 3 times by 1×PBS and a mounting solution (50% glycerol) was used to load and seal the midgut onto a microscope slide for observation.

### 2.7. Fluorescence imaging

A fluorescence microscope (Zeiss Axio Imager.A2, Zeiss HPX120 illuminator) was used for multichannel fluorescence imaging. Hemocytes, nuclei, and F-actin cytoskeletal structures were observed using Rhodamine, DAPI, and FITC filter sets, respectively. Hemocyte behavior was assessed by hemocyte count—the number of granulocytes observed on the dissected midgut by fluorescence microscopy after fixation and CM-Dil staining. Statistical analysis was performed using GraphPad Prism version 10.1.2 (GraphPad Software, Boston, MA, USA).

### 2.8. Mosquito midgut treatment and hemocyte collection for ex vivo analysis

Mosquitoes were cold-anesthetized and injected with 4500 nL of anti-coagulant RPMI (20 mM Sodium citrate, RPMI 1640) to efficiently collect as many hemocytes as possible for the assay. Following an incubation period of 20 mins in RT. The mosquitoes were injected once again with RPMI and the last two abdominal segments were excised and 4.5 µL of liquid was collected by a glass capillary and transferred to a Low Protein Binding Microcentrifuge Tube (Thermo Fisher Scientific, Waltham, MA, USA) on ice. The hemocytes were counted using a cell hemocytometer under light microscopy and titered to 10,000 cells/100 µL. Hemocytes were then stained by mixing 3 parts hemocyte with 1 part Cm-Dil and left to incubate for 20 mins in the dark. The midgut was prepared by isolating the guts in anti-coagulant RPMI to remove existing hemocytes and washed 3 times with RPMI (without anti-coagulant). Organs such as the Malpighian tubules and crop were removed leaving only the midgut. 1 µL of 10% DMSO, PGE_2_, or TXA_2_ (each prepared in 10% DMSO) diluted in RPMI was mixed on a glass slide with 9ul of the stained hemocytes. Dissected midguts were then placed in the mixture and allowed to incubate for 5mins. The midguts were then fixed in 20% formaldehyde for 20 mins. F-actin as well as nuclei were also stained prior to imagining.

### 2.9. Sample preparation and ELISA

An anticoagulant buffer (10 mM EDTA, 30 mM sodium citrate in PBS, pH 8.0) was used to extract hemolymph by pooling 5 mosquitoes per sample. The extractant underwent cleaning by precipitating proteins with an acidified alcohol solution (50 mM HCl in Methanol) which was incubated at -20^°^C for 20-30 mins. Supernatant was collected following centrifugation (14,000 rpm, 10 mins, 4^°^C) and transferred to a low-bind tube. The sample was then dried by SpeedVac (Centrifugal Evaporator CVE-3000, Republic of Korea) for 3 hours at no to minimum heat until a thin film was formed. ELISA buffer was added to reconstitute the sample. We used the Thromboxane B_2_ ELISA Kit (Cayman Chemicals, Ann Arbor, MI, USA) and quantification was performed following manufacturer protocols. The plates were read at 405 nm using a Molecular Devices EMax® Precision Microplate Reader (Molecular devices, San Jose, CA, USA) and a 4 PL parameter logistic plot was used to determine the amount of endogenous TXB_2_ in the mosquito hemolymph based a standard curve obtained from the TXB_2_ standard provided in the kit.

### 2.10. TXA_2_ synthase docking analysis

An atlas of candidate cytochrome P450 PDB files or amino acid sequences were retrieved from UniProt (Bateman et al., 2025), with abbreviations and UniProt accession numbers provided in the Supplementary information (Appendix A). For entries lacking 3-dimensional predictive structures, prediction models were created from the amino acid sequences using AlphaFold2 (DeepMind) with default parameters (Jumper et al., 2021; Varadi et al., 2022). Entries with well-known roles in biosynthesis or detoxification were excluded as were incomplete sequences shorter than 200 amino acids. Entries lacking active site sequence regions were also removed. Using PyMol software (Schrödinger, LLC. (2023). PyMOL Molecular Graphics System, Version 2.5.), the predicted structure of the TBXAS1 (human TXA_2_ synthase, UniProtKB ID: P24557) was aligned with an experimentally determined human CYP21A2 progesterone complex (PDB accession number: pdb_00004y8w) containing a heme molecule. The position of the heme was transferred and merged into human CYP5A1 (TBXAS1) model. Mosquito entries were then spatially aligned with the newly merged human TBXAS1 containing a heme molecule and the degree of similarity between the final mosquito candidate and human model were quantified. We used AutoDockTools 1.5.7. (Morris et al., 2009) to assign Gasteiger charges and merge non-polar hydrogens for both the candidate protein and ligand PGH_2_ (PubChem CID: 445049).

Molecular docking analysis was performed by AutoDock4 with the candidate protein as the target and PGH_2_ as the ligand. Search space was defined using heme coordinates obtained from PyMol alignment.

### 2.11. RNA extraction and RT-qPCR

Whole mosquitoes were homogenized with a pestle in TRIzol®. RNA was extracted by chloroform (Thermo Fisher Scientific, Waltham, MA, USA) and precipitated with isopropanol; pellets formed from centrifugation were washed in 75% Ethanol and resuspended in sterile RNase-free water. RNA was then quantified using Nanodrop (NanoDrop 2000, Thermo Scientific, Waltham, MA, USA). qRT-PCR was performed with 100 ng of total RNA using a TOPreal™ SYBR Green RT-qPCR kit (Enzynomics, Daejeon, Republic of Korea) with gene specific primer sets for TBXAS1 candidates and endogenous controls (Appendix B. Supplementary table1), according to the manufacturer’s instructions. The reaction and measurements were performed using a QuantStudio™ 3 Real-Time PCR System (Thermo Fisher Scientific, Waltham, MA, USA). Relative gene expression was calculated using the 2^−ΔΔ^Ct method. All target genes were normalized using *rl32* as the endogenous control (Field and Smith, 2023). RNA expression levels were calculated by comparing fold change values and statistical analysis was performed.

### 2.12. dsRNA synthesis and RNA interference of candidate genes

Oligonucleotides were designed to specifically amplify target regions for dsRNA synthesis using NCBI Primer-BLAST (https://www.ncbi.nlm.nih.gov/tools/primer-blast/) tool. For in vitro dsRNA production, RT-PCR was first performed on RNA pooled from five female mosquitoes using the SuPrime Script RT-PCR kit (Genetbio, Daejeon, Republic of Korea). Following amplification, T7 ligation was performed using the TaKaRa Ex Taq kit (Takara Bio Inc., Kusatsu, Shiga, Japan) with T7 promoter–containing primers (Appendix B. Supplementary table1). The resulting dsDNA construct was purified using an Expin™ CleanUp SV kit (GeneAll, Seoul, Republic of Korea) and adjusted to a final concentration of 100 ng/µl using a Nanodrop (NanoDrop 2000, Thermo Scientific, Waltham, MA, USA). dsRNA was synthesized using the MEGAscript RNAi Kit (Thermo Fisher Scientific, Waltham, MA, USA) according to the manufacturer’s instructions and diluted to a final concentration of 1000 ng/µl with elution buffer.

Cold-anesthetized female mosquitoes were injected intrathoracically with 100 nL of the dsRNA product. Following a 24 h recovery period, mosquitoes were provided a mixture of 2% SDS in 10% sucrose and allowed to feed *ad libitum* for 24 h, after which an additional 24 h recovery period was given before midgut dissection and downstream analyses.

## 3. Results

### 3.1. Damage signals following blood-feeding and chemical treatment recruit immune cells to the mosquito midgut

The extent and process of damage involving the mosquito midgut is a multi-faceted process that requires further investigation. While natural damage to the midgut occurs during blood feeding events, several factors such as the semi-hematophagous nature of *Culex pipiens* and coagulation of blood within the mosquito midgut when dissected limited the usage of blood feeding as a method of damage induction in experimental settings. Therefore, we used the chemical detergent SDS at 2% concentration mixed with 10% sucrose to induce damage via feeding, as demonstrated in previous studies (Janeh et al., 2017), to mimic the natural damage caused by blood feeding (Fig. 1*A*). A separate SDS feeding assay confirmed sufficient feeding in both the control and SDS-containing sucrose groups, with no statistically significant difference in feeding ratio (Appendix B. Supplementary figure1). Imaging following fluorescence labeling of F-actin, nuclei, and hemocytes showed significant damage to the midgut, as evidenced by disfiguration of the actin structure and misalignment of the actin structural network relative to the nuclei in both SDS-treated and blood-fed midguts (Fig. 1*B*). Hemocyte recruitment increased 13-fold and 7.8-fold following damage by SDS and blood feeding, respectively (Fig. 1*C*). To determine the temporal effect of SDS-induced damage, midguts were dissected and hemocytes attached to the midgut at selected time points after SDS feeding were counted. Hemocyte subpopulations were identified by morphological characteristics (Appendix B. Supplementary figure 2) and CM-DiI (red) fluorescence intensity. Immediately after SDS feeding was discontinued, hemocyte attachment to the midgut increased 3.0-fold compared to controls, rising further to 4.1-fold at 24 hours. By 48 hours post-discontinuation, hemocyte numbers decreased to near-baseline levels, remaining only marginally elevated relative to controls (Fig. 1*D*). The number of total hemocytes and granulocytes also increased by SDS treatment at 2.0- and 1.8-fold (Fig. 1*E-F*), respectively. The proportion of granulocytes relative to the total hemocyte population also increased from 5.2% to 9.3% following treatment (Fig. 1*G*). TXB_2_ levels were approximately two-fold higher in both whole-body and hemolymph samples at 24 h after discontinuation of SDS feeding (Fig. 1*H-I*).

**Figure 1.**
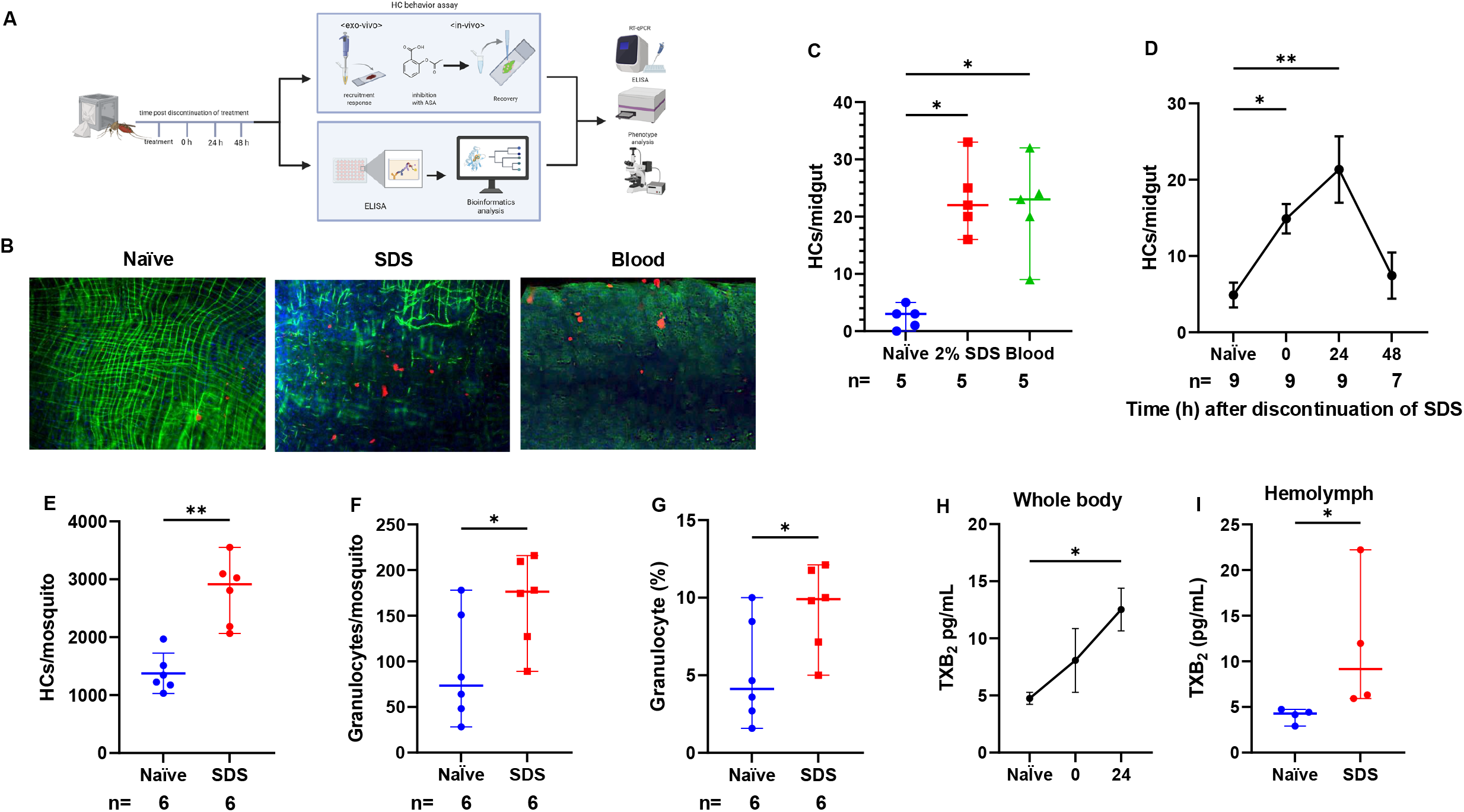
Damage signals recruit immune cells to the mosquito midgut. Sample sizes (n) are indicated below each graph. (a) Overview of the experimental design. (b) Fluorescence images of midgut stained with Phallodin (actin, green), CM-Dil (hemocytes, red), and Hoechst (nuclei, blue). Scale bar indicates 100 µm. (c) Number of hemocytes attached to the midgut under different treatment conditions. Data presented as median ± 95% CI. (d) Time-dependent changes in hemocyte aggregation at the midgut following 2% SDS treatment. Data presented as median ± 95% CI. (e) Number of circulating hemocytes following SDS damage. Data presented as median ± 95% CI. (f) Number of circulating granulocytes following SDS damage. Data presented as median ± 95% CI. (g) Proportion of circulating granulocytes following SDS damage. Data presented as median ± 95% CI. (h) Concentration of TXB_2_ in whole-body extracts following midgut damage. Five biological replicates were analyzed, each consisting of 8 mosquitoes. Data presented as median ± 95% CI. (i) Concentration of TXB_2_ observed in hemolymph following midgut damage. Four biological replicates were analyzed, each consisting of 8 mosquitoes. Statistical comparisons were performed using the Wilcoxon test for pairwise comparisons. For panel (g), differences among groups were assessed using the Kruskal–Wallis test followed by Dunn’s multiple comparisons test. Naïve, sugar feeding stopped after 24 h; Blood, blood feeding stopped after 24 h; SDS, 2% SDS feeding stopped after 24 h. *P < 0.05; **P < 0.01; ***P < 0.001; ****P < 0.0001; NS, P > 0.05.

### 3.2. Roles of TXA_2_ in stimulating hemocyte recruitment to the mosquito midgut

To assess the role of eicosanoid signaling in hemocyte recruitment, acetylsalicylic acid (ASA, aspirin) was administered to inhibit prostanoids production (Fig. 2*A*). Intrathoracic injection of ASA at the time of 2% SDS-induced midgut damage resulted in a 2.9-fold decrease in hemocyte attachment to the midgut after 24 h compared to DMSO-treated controls (Fig. 2*B-C*). Hemolymph extraction and subsequent cell count revealed a decrease in total hemocyte numbers, including granulocytes which showed a decrease from 460 cells to 160 following ASA treatment (Fig.2*D-E*). However, the proportion of granulocytes relative to the total hemocyte population did not differ significantly between control and ASA-treated groups (Fig. 2*F*).

**Figure 2.**
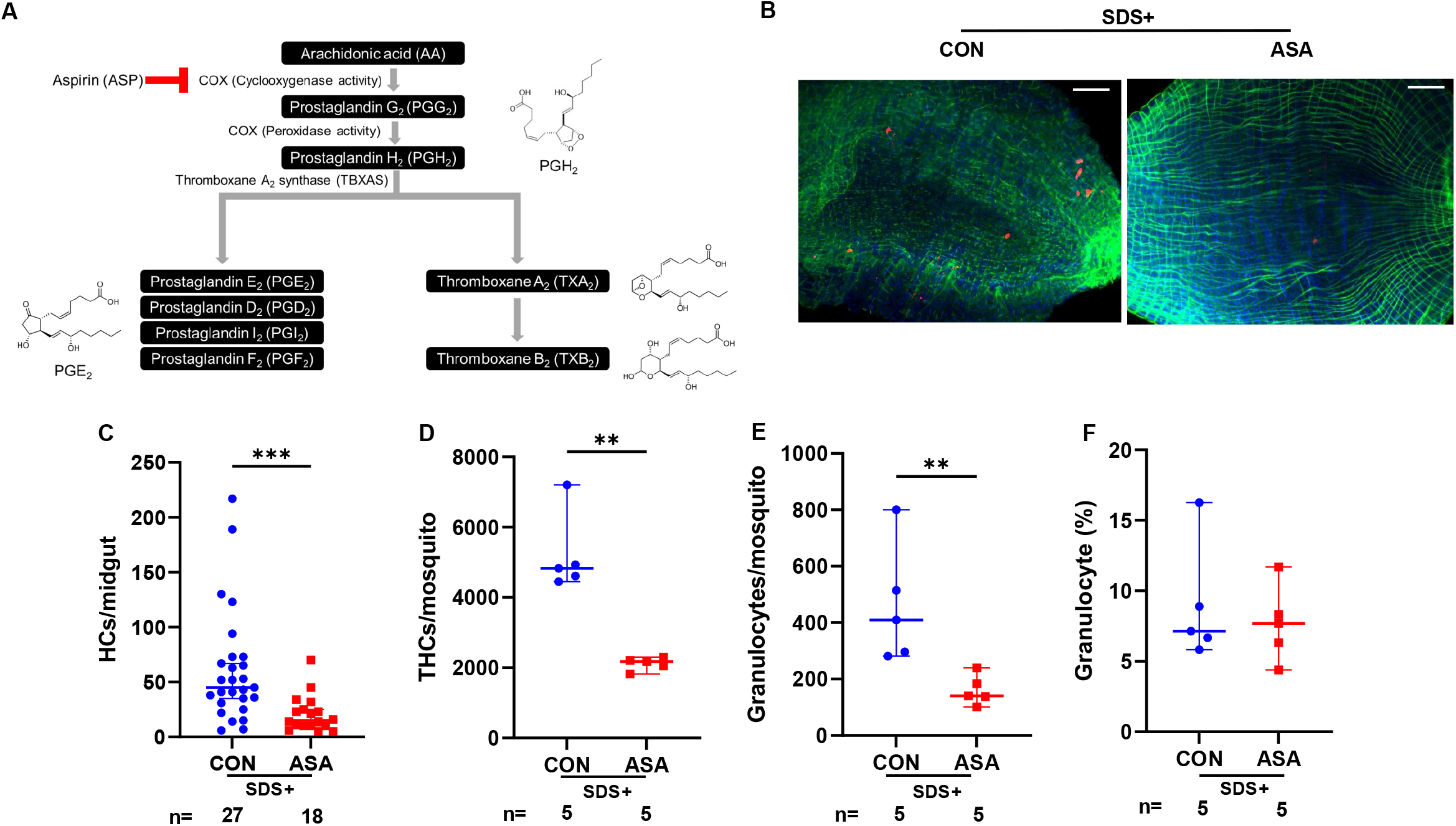
Inhibition of TXA_2_ synthesis by aspirin reduces hemocyte recruitment to the midgut 24 h after 2% SDS treatment. Sample sizes (n) are indicated below each graph. (a) Biosynthetic pathway of prostanoids. Pathways blocked by ASA shown in red. (b) Fluorescence images showing hemocytes attached to the midgut 24 h after SDS treatment. Midgut stained with Phallodin (actin, green), CM-Dil (hemocytes, red), and Hoechst (nuclei, blue). Scale bar indicates 100 µm. (c) Hemocyte recruitment to the midgut inhibited by aspirin (ASA; 100 ng). Data presented as median ± 95% CI. (d) Reduction in the total number of circulating hemocytes following ASA treatment. Data presented as median ± 95% CI. (e) Effect of ASA on the number of circulating granulocytes. Data presented as median ± 95% CI. (f) Changes in granulocyte composition in response to ASA treatment. Data presented as mean ± SE. For panels (d–f), each biological replicate consisted of pooled hemolymph collected from three female mosquitoes. Statistical comparisons were performed using the Wilcoxon rank-sum test. *P < 0.05; **P < 0.01; ***P < 0.001; ****P < 0.0001; NS, P > 0.05. CON, dimethyl sulfoxide (DMSO; 1%, 100 nL per female mosquito); ASA, aspirin (100 ng, 100 nL per female mosquito).

An *ex vivo* assay was performed ahead of time to assess the direct effects of eicosanoids on hemocyte behavior under controlled conditions, thereby minimizing systemic physiological variables present *in vivo*. The use of an anticoagulant helped to dislodge any hemocytes previously attached to organs or injection sites. Exposure of extracted hemocytes and dissected midguts to a dosage of 100 ng TXA_2_ resulted in a 3.0-fold increase in hemocyte recruitment compared with controls (Appendix B. Supplementary figure 3). In contrast, PGE_2_ treatment did not produce a statistically significant increase in recruitment.

We then examined the influence of TXA_2_ on hemocyte behavior *in vivo*. Increasing doses of TXA_2_ resulted in a steady rise in hemocyte attachment to the midgut, with a 3.2-fold increase observed at 3 ng and a 5.4-fold increase at 10 ng (Fig. 3*A-B*) compared to controls. Analysis of the hemolymph following a 10 ng TXA_2_ treatment revealed a 1.8-fold increase in total circulating hemocytes, indicating an activated immune state (Fig. 3*C*). Although the absolute number of granulocytes in the hemolymph remained relatively stable in the treatment group (Fig. 3*D*), their proportional representation decreased from an average of 9% to 5% following TXA_2_ injection (Fig. 3*E*).

**Figure 3.**
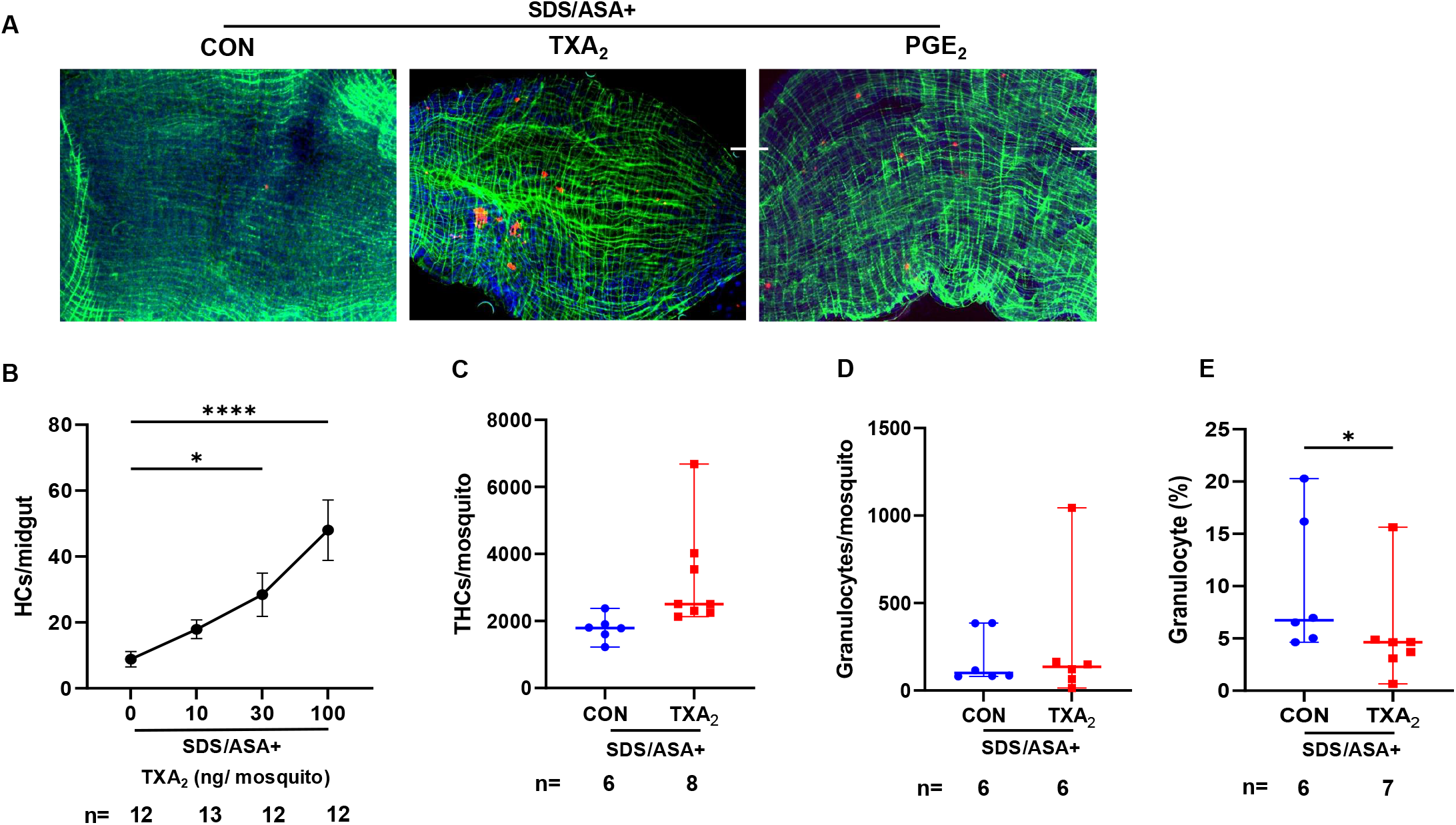
TXA_2_ promotes hemocyte recruitment (recovery) to the mosquito midgut following aspirin treatment. Sample sizes (n) are indicated below each graph. (a) Fluorescence images of the midgut showing the rescue effects of TXA_2_. Midgut stained with Phallodin (actin, green), CM-Dil (hemocytes, red), and Hoechst (nuclei, blue). Scale bar indicates 100 µm. (b) Dosage-dependent effect of TXA_2_ on hemocyte recruitment. Data presented as median ± 95% CI. (c) Effect of TXA_2_ treatment on the total number of circulating hemocytes during recovery. Data presented as median ± 95% CI. (d) Effect of TXA_2_ treatment on the number of circulating granulocytes during recovery. Data presented as median ± 95% CI. (e) Changes in the proportion of circulating granulocytes following TXA_2_ treatment. Data presented as median ± 95% CI. For panels (c–e), each biological replicate consisted of pooled hemolymph collected from three female mosquitoes. Panel (b) was analyzed using the Kruskal–Wallis test followed by Dunn’s multiple comparisons test. Panels (c–e) were analyzed using the Wilcoxon rank-sum test. P < 0.05; P < 0.01; P < 0.001; P < 0.0001; NS, P > 0.05. CON, dimethyl sulfoxide (DMSO; 1%, 100 nL per female mosquito); TXA_2_, TXA_2_ (10 ng, 100 nL per female mosquito).

### 3.3. Structural and biochemical evidence for TXA_2_ production in mosquitoes

Phylogenetic analysis of CYP enzymes did not clearly resolve candidate thromboxane A_2_ synthases. To identify potential functional candidates, molecular docking simulations were performed using PGH_2_ as the precursor for TXA_2_ (Fig. 4*A*). Candidates were evaluated based on predicted binding affinity and structural similarity to the human thromboxane A_2_ synthase CYP5A1, also known as TBXAS1 (Fig. 4*B-C*). Following removal of duplicated, misannotated, or truncated entries, 45 sequences were retained for analysis. Of these, 18 had predicted structures available, while structural models for the remaining 27 were generated using AlphaFold. Docking of PGH_2_ was conducted using AutoDock4 via AMDock. Using a threshold of predicted binding affinity less than −6 kcal/mol, 15 candidates were retained for further evaluation. Candidates were subsequently prioritized based on predicted binding affinity together with structural plausibility, including ligand orientation and complementarity to the active site geometry. Through this filtering process, five candidates were selected for sequencing. Based on the experimentally obtained sequence data, removal of redundant and non-expressed proteins resulted in CYP4F14 and CYP6D3 as the final candidates. Structural comparison with human TBXAS1 showed that while both candidates exhibited similar overall charge distributions, only CYP6D3 displayed a similar hydrophobic substrate-binding environment to CYP5A1 and achieved an active site root-mean-square deviation value below 1 Å in structural alignment. Docking analysis further revealed that in CYP6D3, the dominant docking cluster corresponded to the lowest-energy pose, positioning the PGH_2_ endoperoxide bridge above the heme iron in a catalytically competent orientation, whereas this geometry was not consistently observed for CYP4F14. In CYP6D3, the oxygen atom proximal to C9 was positioned 3.4 Å from the heme iron, consistent with a mechanistically plausible range for catalytic turnover and comparable to the 2.9 Å shown in the human model (Appendix B. Supplementary figure 4). It was also shown to have a binding energy of −8.23 kcal/mol, comparable to that of the human model (−7.93 kcal/mol). Residues within 4 Å of the docked PGH_2_ included His119, Phe121, Asn122, Phe213, Arg217, Val300, Leu303, Ala304, Glu307, Thr308, Val370, Phe484, and Val485, consistent with a predominantly hydrophobic substrate-binding pocket with polar residues likely contributing to ligand positioning (Fig. 4*D-E*).

**Figure 4.**
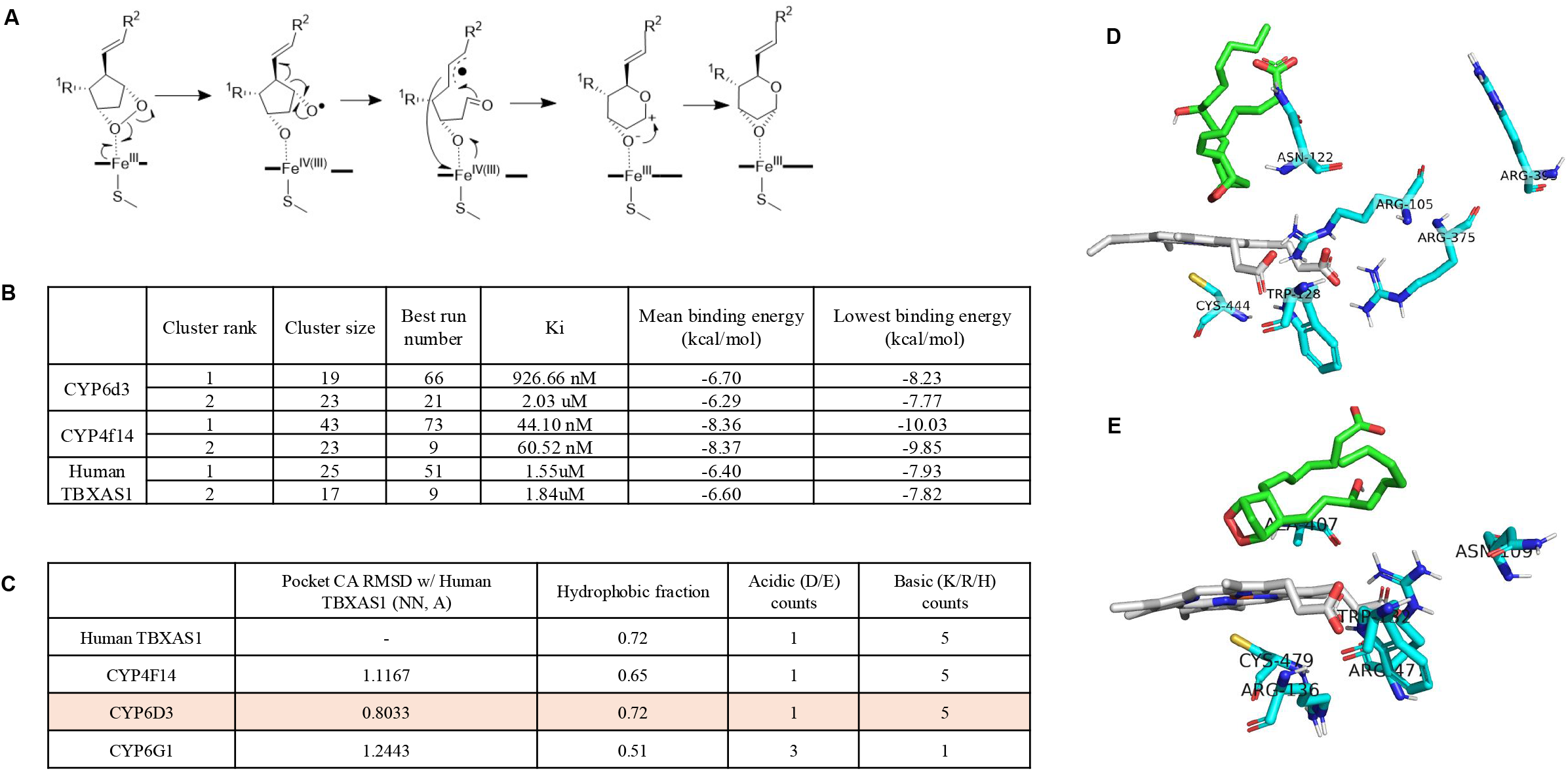
Structural and evidence of TXA_2_ production. (a) Proposed mechanism of TXA_2_ synthesis. (b) CYP6D3 shows a pocket CA RMSD (Pocket C-alpha Root Mean Square Deviation) <1 indicating strong similarity to CYP5A1 (human) as well as the same hydrophobic fraction and acid, base count. (c) Lowest binding energy of 100 molecular docking runs show similar results for CYP6D3 and CYP5A1 (Human TBXAS1). (d) Active site residues and docked PGH_2_ ligand of CYP6D3 (e) Active site residues and docked PGH_2_ ligan d of CYP5A. Pocket CA RMSD (root-mean-square deviation of aligned C_α_ atoms).

Additionally, while the CYP4F14 model showed a shorter Fe–O distance (2.9 Å) and lower binding energy (−10.03 kcal/mol), its active site architecture and residue composition diverge substantially from that of human TBXAS1, resulting in ligand orientations less consistent with a catalytically relevant configuration compared to CYP6D3.

### 3.4. RNAi Knockdown of TXBXAS1 prevents hemocyte recruitment to the midgut

Knockdown of candidate and control genes was conducted to experimentally validate the functional roles of TXA_2_ in the mosquito midgut damage response (Fig. 5*A-B*). RT-qPCR of the mRNAs coding for *CYP6D3* and *CYP4F14* was performed following SDS damage induction. Relative mRNA levels normalized to *rl32* showed a 2-fold increase at 24 hours after SDS discontinuation in *CYP6D3*, whereas no significant change was observed in *CYP4F14* (Appendix B. Supplementary figure 5A). RNA expression studies of *CYP6D3* in the hemolymph showed a 35-fold change at 24 h post discontinuation of SDS compared to naïve (Fig. 5*C*). RT-qPCR conducted following injections of dsRNA showed a 75-fold decrease in *CYP6D3* expression levels, and a 30-fold decrease in *CYP4F14* expression levels, pointing to significant inhibition of mRNA expression (Fig. 5*D*). Following verification of RNAi knockdown, the phenotype resulting from RNAi showed a 4-fold decrease in hemocyte recruitment to the midgut in both dsCYP6D3- and dsCYP4F14-treated mosquitoes (Fig. 5*E*). ELISA was also performed to quantify TXB_2_ levels following dsRNA injections. The results showed a 25% decline of TXB_2_ levels in dsCYP6D3-treated samples, while TXB_2_ levels for the dsLacZ- and dsCYP4F14-treated samples remained at expected levels (Fig. 5*F*).

**Figure 5.**
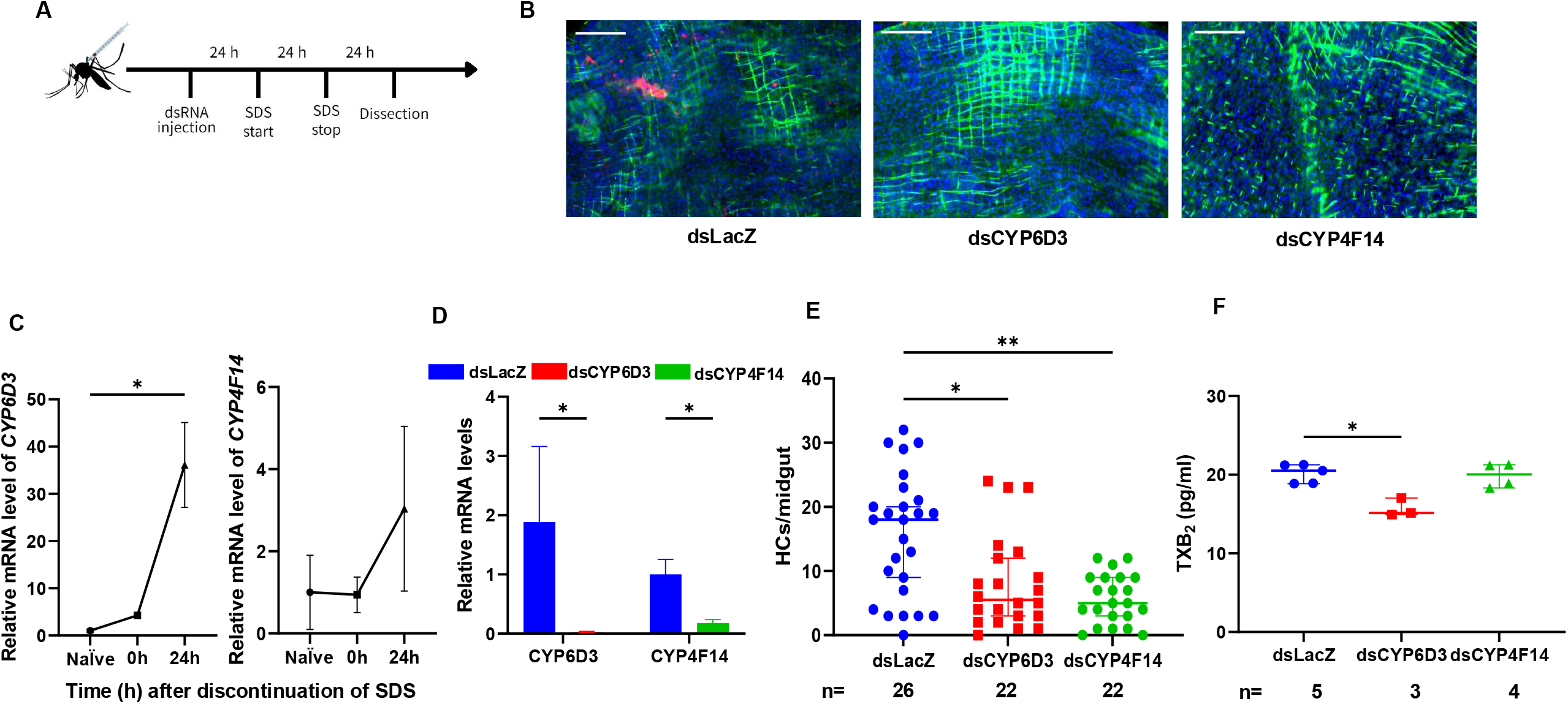
Silencing of TBXAS1 homologs in *Cx. pipiens molestus* inhibits hemocyte recruitment to the midgut. (a) Experimental timeline of RNAi treatment. (b) Fluorescence images of the midgut following RNAi treatment. Midgut stained with Phallodin (actin, green), CM-Dil (hemocytes, red), and Hoechst (nuclei, blue). (c) mRNA expression levels of respective CYP genes measured in the hemolymph. Data are presented as the mean ± SE. (d) mRNA expression levels of respective CYP genes following RNAi treatment. Data presented as median ± 95% CI. (e) Number of hemocytes recruited to the midgut following RNAi treatment. Data presented as median ± 95% CI. (f) TXB_2_ concentration following RNAi treatment. Data are presented as the median ± 95% CI. Five (dsLacZ), three (dsCYP6D3), and four (dsCYP4F14) biological replicates were analyzed, each consisting of five mosquitoes. Panel (d) was analyzed using the Wilcoxon rank-sum test. The remaining panels were analyzed using the Kruskal– Wallis test followed by Dunn’s multiple comparisons test. dsLacZ, double-stranded *LacZ*-injected control; dsCYP6D3, double-stranded *CYP6D3*-injected group; dsCYP4F14, double-stranded *CYP4F14*-injected group. *P* < 0.05; **P** < 0.01; ***P*** < 0.001; **P** < 0.0001; NS, *P* > 0.05.

## 4. Discussion

Despite the global burden of mosquito-borne diseases on human health, mechanisms governing pathogen transmission within mosquitoes remain poorly understood. In particular, early hemocyte aggregation, a key process limiting pathogen dissemination, is not well characterized. Hemocytes are recruited to sites of midgut damage via recognition of damage- and pathogen-associated molecular patterns. Under homeostatic conditions, hemocytes are sessile tissue-bound, residing on structures such as the epidermis and fat body. Upon infection or injury, they become activated and localize to sites of damage. Here, lipid mediators, including eicosanoids, serve as key regulators of the mosquito immune response (Barletta et al., 2019).

Hemocytes are thought to attach to the midgut following damage induced by blood feeding, representing an early step in the recovery process, as suggested by studies examining post-attachment responses (Cardoso-Jaime and Dimopoulos, 2025; Hillyer and Strand, 2014). SDS feeding reliably induces midgut damage and hemocyte recruitment, supporting its use as a reproducible model to study injury-associated immune responses. Analysis of recruitment dynamics revealed an increased proportion of granulocytes following damage, consistent with their role as the primary immune effector cell type in mosquitoes. Granulocytes mediate defense at the midgut through adhesion to the basal lamina, phagocytosis of invading pathogens, and secretion of immune effectors, indicating that their enrichment reflects an active response to tissue damage (Taracena et al., 2018; Hall et al., 2025).

Furthermore, the reduction in hemocyte recruitment following ASA treatment suggests a functional role for eicosanoid signaling in mediating immune cell responses to midgut damage. In mammals, COX catalyzes the conversion of arachidonic acid into prostaglandin H_2_ (PGH_2_), a key precursor for downstream prostanoids including PGE_2_ and TXA_2_. ASA irreversibly inhibits COX activity, thereby limiting the availability of PGH_2_ for prostanoid synthesis. Although canonical COX enzymes have not been identified in insects, COX-like or peroxidase-mediated pathways have been proposed to generate prostaglandin or thromboxane (Fig. 2*A*). Within this context, the decrease in hemocyte recruitment observed here supports a conserved role for eicosanoid-derived signals in regulating immune cell behavior. Notably, the reduction in total hemocyte numbers without a shift in granulocyte proportion suggests that ASA does not selectively affect specific hemocyte subtypes, but rather broadly suppresses hemocyte mobilization or survival (Fig. 2*D-F*). In Anopheles gambiae, PGE2 enhances hemocyte responsiveness and chemotactic activity and has been associated with increased granulocyte representation during immune challenge, suggesting that lipid mediators play a key role in regulating hemocyte dynamics (Ramirez et al., 2015). Together, these findings are consistent with a model in which prostanoid signaling—potentially via TXA_2_—contributes to hemocyte recruitment following tissue damage (Fig. 3). The differential effects of TXA_2_ and PGE_2_ on hemocyte recruitment suggest distinct functional roles for these eicosanoids in mosquito immunity. While PGE_2_ has been previously associated with immune priming, the robust increase in hemocyte recruitment observed following TXA_2_ exposure supports a role for TXA_2_ as a damage-associated signal that actively promotes hemocyte trafficking to the midgut. In addition to this, the decrease in granulocyte proportion within the hemolymph following TXA_2_ treatment, despite relatively stable absolute numbers, is consistent with redistribution of hemocytes from circulation to the midgut. Together, these findings support a model in which TXA_2_ signaling facilitates targeted hemocyte recruitment to sites of tissue damage. Although CYP4F14 exhibited stronger predicted binding affinity to PGH_2_, its docking poses did not consistently position the endoperoxide moiety in a catalytically competent orientation relative to the heme iron. In contrast, CYP6D3 displayed binding energies comparable to human TBXAS1 and consistently positioned the substrate in a geometry compatible with catalytic turnover (Fig. 4*D-E*). Together with its structural similarity to human TBXAS1, these results support CYP6D3 as a more likely functional analog of TBXAS1 in mosquitoes.

Our combined structural, biochemical, and functional analyses identify CYP6D3 as a strong candidate enzyme mediating TXA_2_ production in mosquitoes. Molecular docking and structural comparisons revealed that CYP6D3 closely resembles human TBXAS1 (Yanai and Mori, 2011) with favorable ligand orientation and active-site compatibility for PGH_2_ binding. Consistent with this, RNAi-mediated knockdown of *CYP6D3* resulted in reduced hemocyte recruitment, supporting a functional role in the TXA_2_-dependent damage response pathway. Although ELISA-based quantification of TXB_2_ showed only modest changes, the observed trend was consistent with reduced thromboxane B_2_ production following *CYP6D3* inhibition. Given the functional diversity and redundancy of CYPs, as well as their often promiscuous interactions with eicosanoid substrates—as illustrated by other studies pertaining to the Spodoptera TXA_2_ synthase (Al Baki et al., 2021) —we cannot exclude the possibility that additional *CYP*s contribute to TXA_2_ biosynthesis. In this context, *CYP4F14* may function in a downstream or parallel pathway influencing hemocyte recruitment, although its precise role remains less clearly defined. Taken together, our findings identify TXA_2_-mediated signaling as a key regulator of hemocyte recruitment to the mosquito midgut following damage induction, supporting the presence of a functional, albeit non-canonical, eicosanoid pathway in mosquitoes. By integrating functional assays with molecular docking and comparative structural analysis, we provide evidence implicating specific *CYP* as candidate, *CYP6D3*, mediators of TXA_2_ biosynthesis. Although the precise enzymatic steps and intermediate metabolites remain to be fully resolved, and species-specific differences may limit direct generalization beyond *Culex spp*, our results establish a mechanistic framework linking eicosanoid signaling to early immune responses at the midgut interface. This work not only advances our understanding of mosquito innate immunity but also highlights previously uncharacterized components of eicosanoid metabolism as potential targets for disrupting pathogen transmission.

## CRediT authorship contribution statement

**Dawon Lee**: Conceptualization, Investigation, Formal analysis, Methodology, Software, Data curation, Visualization, Writing – original draft, Writing - Review & Editing. **Du-Yeol Choi**: Conceptualization, Investigation, Formal analysis, Methodology, Data curation, Writing – original draft, Writing - Review & Editing. ***Ju Yeong Kim***: Conceptualization, Resources, Supervision, Project administration, Funding acquisition, Writing – review & editing. **Yun Soo Jang**: Investigation. **Dongjun Kang, Jun Ho Choi**: Visualization, Formal analysis. **Singeun Oh, Arwa Shatta, Myung-hee Yi, Chaerin Park, In-Yong Lee**: Writing – review & editing.

## Conflict of interest

The authors declare no conflict of interest.

**Appendix A. Supplementary data**

**Appendix B. Supplementary data**

## Data availability

All data will be made available upon request.

## Declaration of generative AI and AI-assisted technologies in the manuscript preparation process

During the preparation of this work, the authors used ChatGPT/OpenAI and Gemini/Google to review and edit code used in the molecular docking process and to perform spelling and language checks. After using these tools, the authors reviewed and edited the content as needed and take full responsibility for the content of the published article.

---

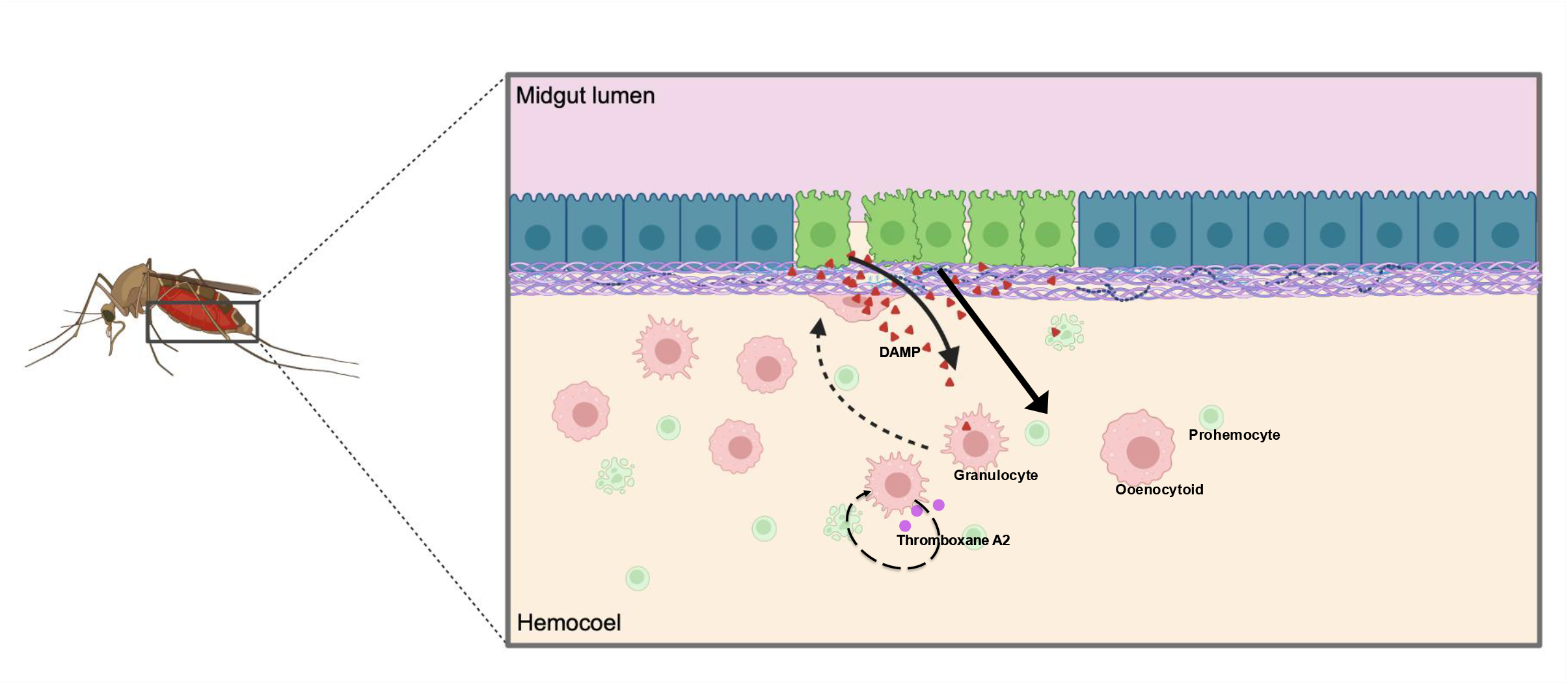

