## Appendix B for "Midgut damage triggers thromboxane A_2_–dependent hemocyte recruitment in *Culex pipiens molestus*"

**Supplementary Table1.** Primers used for sequencing.

| Target | Primer | sequence |
| --- | --- | --- |
| *6d3* | Cx_P450_6d3_2_F | CGT CAG CAC GTG AGA GAA GTC |
|  | Cx_P450_6d3_2_R | GCA CTT CTC CCC GAT CGA AAA |
| *4f14* | Cx_P450_4F14_2_F | TCT TCC TAC TTT CGT GGA TCG C |
|  | Cx_P450_4F14_2_R | CAA GAT CGA AGG ACT GAA ACA CA |
| *lacZ* | T7+dsLacZ_F | TAA TAC GAC TCA CTA TAG GGG AGT CAG TGA GCG AGG AAG C |
|  | T7+dsLacZ_R | TAA TAC GAC TCA CTA TAG GGT ATC CGC TCA CAA TTC CAC A |
| *6d3* | T7+6d3_F2 | TAA TAC GAC TCA CTA TAG GGC GTC AGC ACG TGA GAG AAG TC |
|  | T7+6d3_R2 | TAA TAC GAC TCA CTA TAG GGG CAC TTC TCC CCG ATC GAA AA |
| *4f14* | T7+4f14_F2 | TAA TAC GAC TCA CTA TAG GGT CTT CCT ACT TTC GTG GAT CGC |
|  | T7+4f14_R2 | TAA TAC GAC TCA CTA TAG GGC AAG ATC GAA GGA CTG AAA CAC A |
| *rl32* | RL32_F | AAG CCG AAA GGT ATC GAC AA |
|  | RL32_R | CAG TAG ACG CGG TTC TGC AT |
| *6d3* | cx_6d3_q4F | GAC GAT TGT GGC TGC CGG GAA |
|  | cx_6d3_q4R | GTC GTG TAC CTC GCC AGC AGA T |
| *4f14* | cx_4f14_q4F | GGC TAT CGG TAC GCG TGG CT |
|  | cx_4f14_q4R | CTG GCA ACC CTG CGC AAT CC |
| *6g1* | 6g1_1F | CGC ACC TGC CTG GGG TCT C |
|  | 6g1_1R | TCG AGC ACA GCG TGA AGG CA |

Supplementary Figure S1. Feeding ratio following ingestion of a 7% sucrose solution. Feeding ratio was defined as the proportion of mosquitoes with visibly blue abdomens relative to the total experimental population. Results are based on three independent biological replicates (total n = 84 mosquitoes; CON, n = 45; SDS, n = 39).

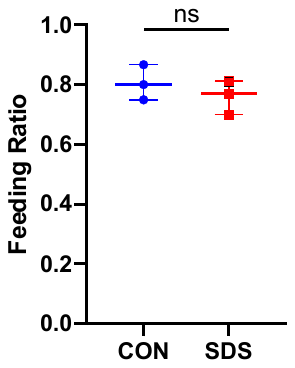

Supplementary figure 2. Morphological characterization of hemocytes. The hemocytes was observed under the steric microscope with 400× magnification. Scale bar indicates 10 µm.

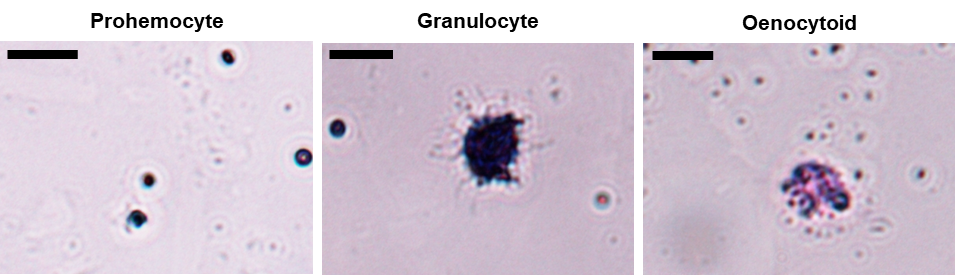

**Supplementary figure 3.** Exo vivo experiment shows the number of hemocytes attached to the midgut following under exposure to 1000 ng of respective chemicals. CON, 1% dimethyl sulfoxide (DMSO; 100 nL per female mosquito); TXA₂, TXA₂ (10 ng, 100 nL per female mosquito). *P < 0.05; **P < 0.01; ***P < 0.001; ****P < 0.0001; NS, P > 0.05.

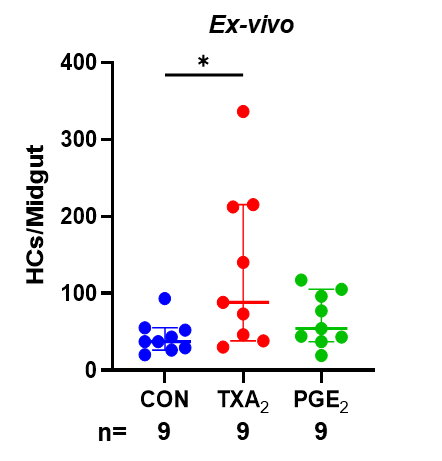

**Supplementary figure 4.** Structural results of CYP450 docking with PGH_2_ (a) CYP4F14: Coordinate of central grid of maps: (-56.339, 15.427, 26.787), distance between Heme iron and closest oxygen atom: 2.7 Å (b) CYP6D3 Coordinate of central grid of maps: (-36.154, 49.923, 27.574), distance between Heme iron and closest oxygen atom: 3.4 Å, (c)Human CYP5A1: Coordinates of Central Grid Point of Maps = (-56.107, 15.661, 27.106), distance between Heme iron and closest oxygen atom: 4.0 Å

A

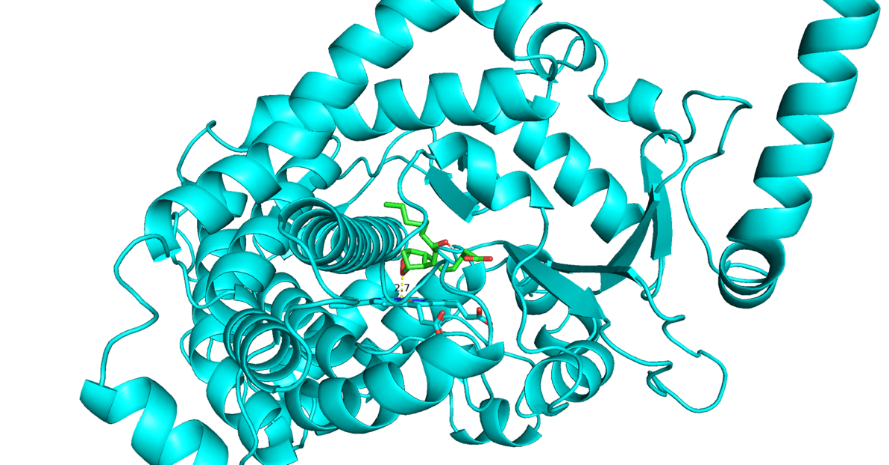

B

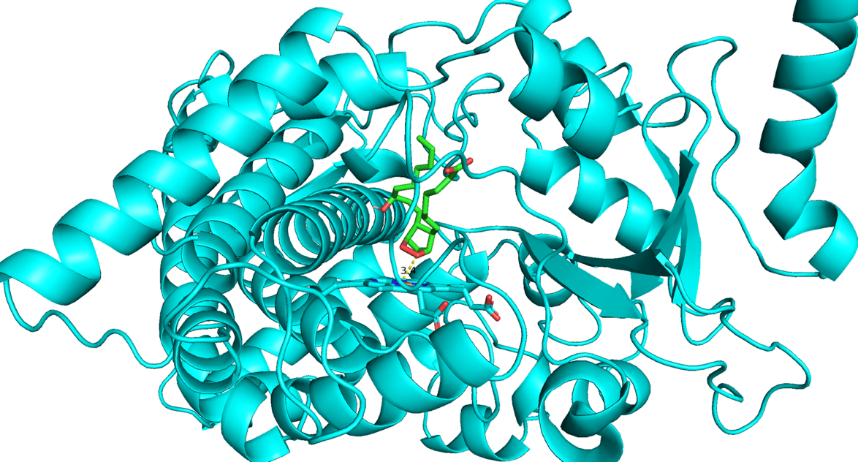

C

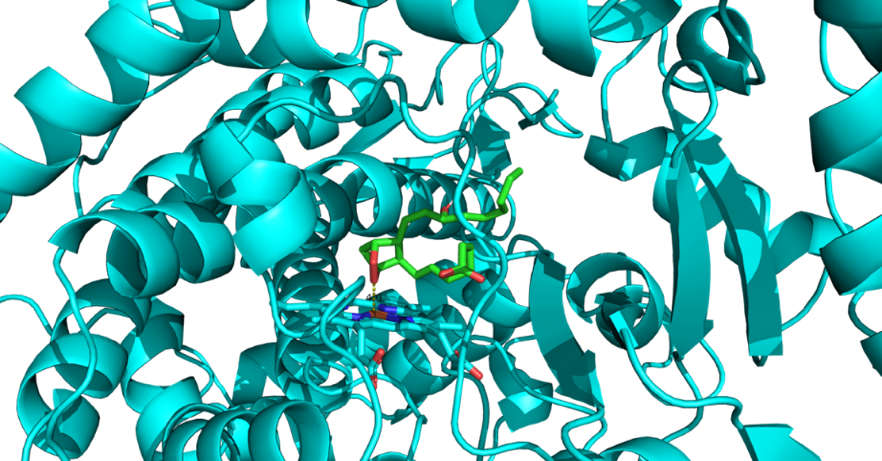

**Supplementary figure 5.** (a) RNA expression profiles of candidate TBXAS1 homologs in the whole body. (b) RNA expression profiles of candidate TBXAS1 homologs in the gut. HC, hemocytes. *P < 0.05; **P < 0.01; ***P < 0.001; ****P < 0.0001; NS, P > 0.05.

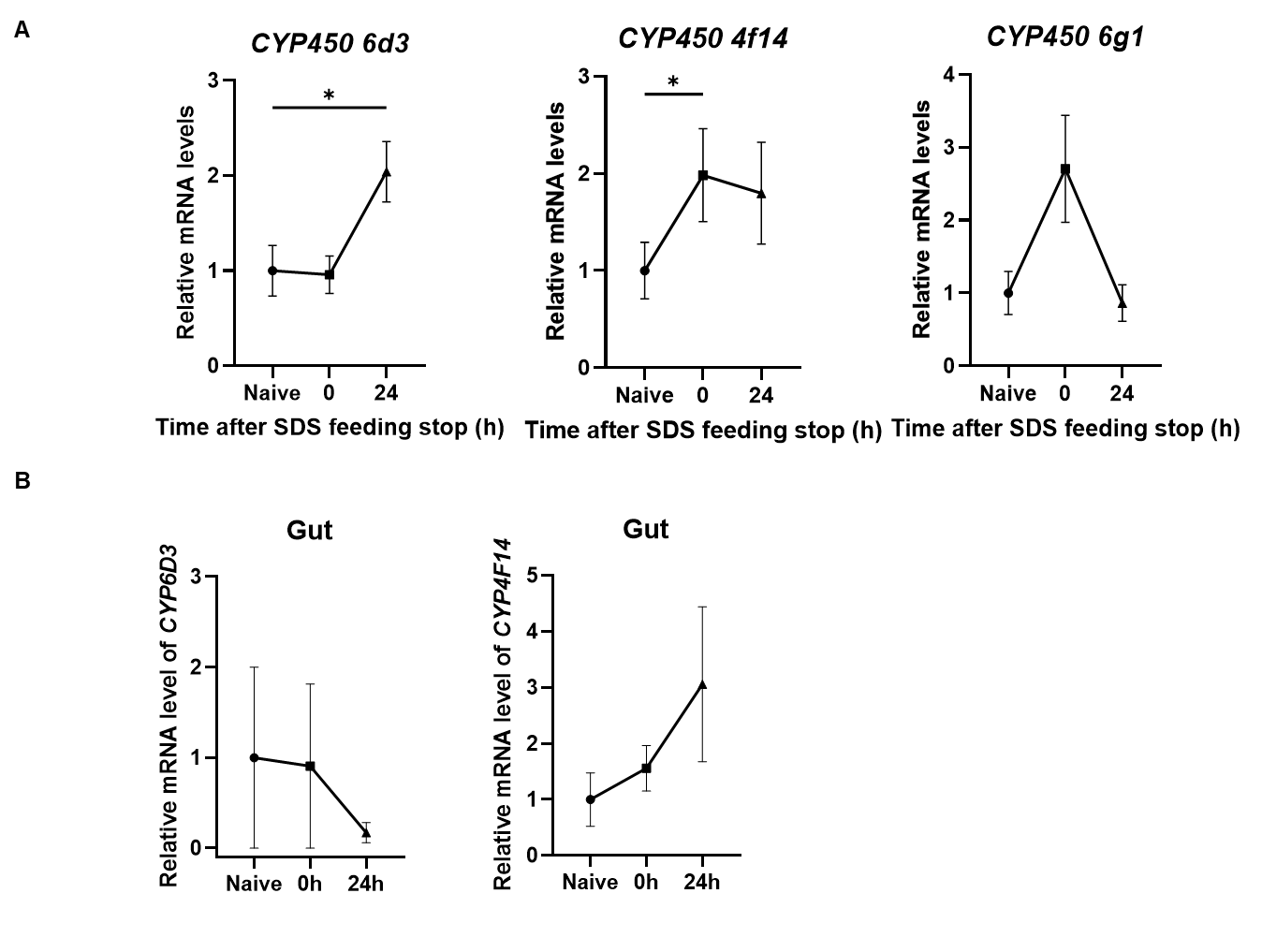
